# Trophic eggs affect caste determination in the ant *Pogonomyrmex rugosus*

**DOI:** 10.1101/2023.01.28.525977

**Authors:** E. Genzoni, T. Schwander, L. Keller

## Abstract

Understanding how a single genome creates distinct phenotypes remains a fundamental challenge for biologists. Social insects provide a striking example of polyphenism, with queen and worker castes exhibiting morphological, behavioural, and reproductive differences. Here we show that trophic eggs, which do not contain an embryo and are primarily regarded as a source of food, play a role in the process of caste determination in the harvester ant *Pogonomyrmex rugosus*. When first instar larvae were given access to trophic eggs, they mostly developed into workers. By contrast, larvae without access to trophic eggs developed into queens. We found that trophic eggs differ in many ways from viable eggs, including texture, morphology and their contents of protein, triglycerides, glycogen, sugar and small RNAs. Moreover, comparison of miRNA fragment size distributions suggests differences in the composition of miRNAs between the two egg types. This is the first demonstration of trophic eggs playing a role in caste determination in social insects.

## Introduction

Many species of insects, spiders, amphibians, marine invertebrates and sharks produce trophic eggs, a special type of eggs that do not contain an embryo (Levin and Bridges 1995; Blake and Arnofsky 1999; Collin 2004; Kudo and Nakahira 2004; Perry and Roitberg 2006; Strathmann and Strathmann 2006; Gibson *et al*. 2012; López-Ortega and Williams 2018). It is generally assumed that these non-developing eggs are either a by-product of failed reproduction or that they serve as nutrition for offspring (Perry and Roitberg 2006). However, the suggestion that trophic eggs solely provide a nutritional function is based on surprisingly little evidence. We here report a direct function of trophic eggs in the determination of alternative phenotypes in ants.

Trophic eggs have been reported in many ant species (Figure 1 and Supplementary Table 1) and are generally thought to mostly or only serve as food for offspring (Crespi 1977; Hölldobler and Wilson 1990). Trophic eggs play an important role during the time of colony founding (Hölldobler and Wilson 1990). Because in most ant species queens do not forage after the mating flight, they metabolize the alary muscles and fat bodies and convert them into eggs, which serve as food to rear the first batch of larvae (Huber 1905; Keller and Passera 1989). Except for a few species such as *Crematogaster smithi (*Heinze *et al*. 1995) and *Acanthomyrmex ferox* (Gobin and Ito 2000), the queens generally stop producing trophic eggs after the eclosion of the first workers (see Figure 1 and Supplementary Table 1). So far the absence of trophic eggs has been reported in only one species (Supplementary Table 1).

**Figure 1.**
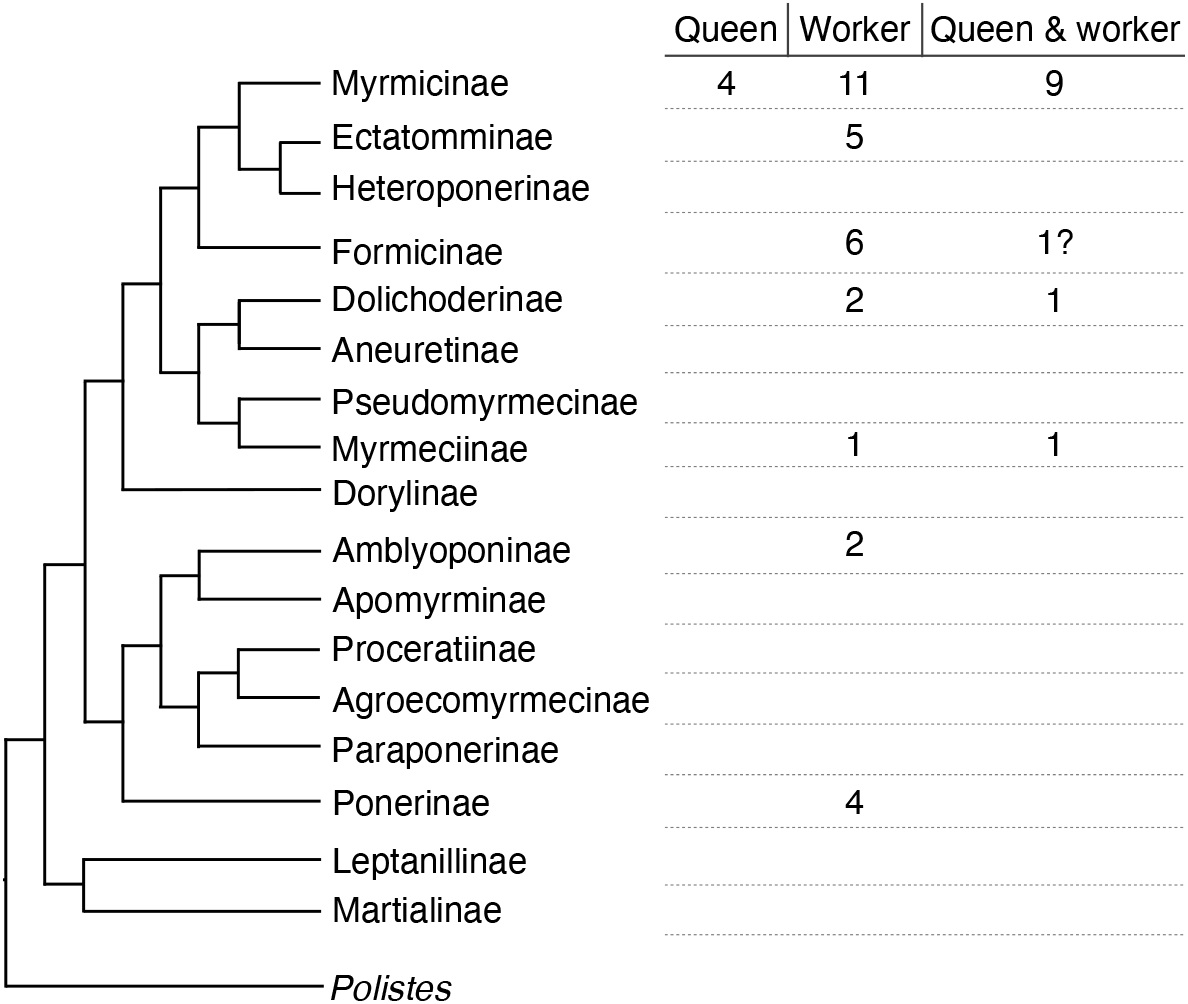
Trophic egg production is widespread in ants. Simplified phylogenetic tree of ant subfamilies redrawn after Romiguier *et al*. (2022). The number of species with documented trophic egg production by queens, workers or both castes is indicated for each subfamily. The question mark indicates that it is unclear whether trophic eggs can be produced by queens (in *Lasius niger* trophic eggs are produced by workers and possibly queens, see supplementary Table 1). Details on the species and related references can be found in Supplementary Table 1.

In many ant species, workers are capable of producing haploid males but lack a spermatheca and the ability to produce diploid female offspring (Hölldobler and Wilson 1990; Bourke 1988; Hammond and Keller 2004; Wenseleers and Ratnieks 2006). In some species, workers further lay trophic eggs (see supplementary Table 1). These eggs have a trophic function, in particular in genera such as *Pogonomyrmex* where there is no or only little trophallaxis (regurgitative food sharing) among workers. In such species it is has been suggested that trophic eggs may play an important role in food distribution within the colony (Gobin and Ito 2000).

Several studies suggested that the presence of trophic eggs may affect the developmental trajectory of larvae. A study in the Argentine ant *Linepithema humile* showed that the presence of queens in colonies was associated with a drastic decrease in the number of worker-laid trophic eggs as well as a decrease in the proportion of larvae developing into queens (Bartels 1988). Bartels thus proposed that trophic eggs may increase the probability of larvae to develop into queens, which are much larger than workers (Bartels 1988). In some lineages of *Pogonomyrmex barbatus*, queens laid a higher proportion of trophic eggs upon the experimental increase of maternal juvenile hormone (Helms Cahan *et al*. 2011). Because the treatment also strongly reduced the number of workers produced but triggered a 50% increase in worker body size, trophic eggs were suggested to affect worker size. An increase in the proportion of trophic eggs has also been suggested to be associated with an increase in the proportion of larvae developing into queens (*L. humile*: Bartels 1988; *P. barbatus*: Helms Cahan *et al*. 2011). Finally, because they observed an increase in nitrogen content with increasing female caste size in the ant *Pogonomyrmex badius*, Smith and Suarez (2010) suggested that the larger castes may consume more trophic (nutritional) eggs than the smaller caste which would feed more on foraged insects. In summary, in at least three species of ants, trophic eggs may play a role in the developmental trajectories of female larvae.

While conducting egg cross fostering experiments in the ant *P. rugosus* to study worker size variation, we observed a sudden increase in the frequency of females developing into queens. During these experiments, we discarded trophic eggs and only cross fostered eggs that would eventually hatch (hereafter viable eggs). This raises the possibility that the absence of trophic eggs influenced the process of caste determination. These observations prompted us to investigate whether trophic eggs play a role in caste determination in *P. rugosus*. Our experiments revealed that the presence of trophic eggs reduces the probability that female larvae develop into reproductive individuals. Metabolomic analyses also revealed profound differences between viable and trophic eggs, including in the composition of miRNAs and content of protein, triglycerides, glycogen, and sugar.

## Materials and methods

Some populations of *Pogonomyrmex rugosus* are characterized by a genetic caste determination system whereby development into queens or workers is determined by whether the eggs are fertilized by males of the same genetic lineage or a different genetic lineage than the queen producing the eggs (Helms Cahan *et al*. 2002; Julian *et al*. 2002; Volny and Gordon, 2002; Helms Cahan and Keller 2003). We collected queens from two populations (Bowie and Florence, Arizona, USA) known to harbor only colonies with non-genetic caste determination (Schwander *et al*. 2007). The queens were collected after mating flights in 2008 (Bowie) and 2023 (Bowie and Florence) to initiate colonies in the laboratory. The colonies were maintained in plastic boxes containing water tubes (glass tubes filled with water and sealed with a cotton plug) at 28°C and 60% humidity, with a 12-h/12-h light:dark cycle. They were fed *ad libitum* once a week with grass seeds, flies and 20% honey water. Eggs were collected in October 2020 for the experiment investigating the effect of trophic eggs on larval caste fate, in November 2021 for estimating the percentage of trophic eggs and from February to December 2021 for the egg content analyses.

All statistical analyses were performed with Rstudio (RStudio Team 2015).

### Trophic and viable egg production

To verify that workers do not lay trophic eggs, as previously shown for other *Pogonomyrmex* species (Supplementary Table 1), we created 12 queenless colonies (by removing the queen) and waited approximately three weeks until workers started laying eggs. From each of these colonies, we isolated two groups of five workers for 12 hours every two days for two weeks in November 2020 to obtain eggs. Collected eggs were then placed for 10 days in a petri dish containing a water reservoir to study their development and distinguish whether they were trophic or viable.

To determine whether queens laid variable percentages of trophic eggs over time, we isolated each of 43 *P. rugosus* queens for 8 hours every day for 2 weeks, before and after hibernation, and counted the number of trophic and viable eggs laid (see results for how to discriminate the two types of eggs). We hibernated the queens because we previously showed that hibernation is important to trigger the production of gynes in *P. rugosus* colonies in the laboratory (Schwander *et al*. 2008; Libbrecht *et al*. 2013). Hibernation conditions were as described in Libbrecht *et al*. (2013). The percentage of trophic eggs was compared using a linear mixed effect model with before vs after hibernation as the explanatory variable and colony as a random factor.

To assess whether viable and trophic eggs were laid in a random order, or whether eggs of a given type were laid in clusters, we isolated 11 queens for 10 hours, eight times over three weeks, and collected every hour the eggs laid. To determine whether viable and trophic eggs were laid in a random order, we performed a Wald–Wolfowitz runs test for each queen’s egg laying sequence (package *snpar* v.1.0; this non-parametric test calculates the likelihood that a binomial data sequence is random).

### Trophic egg influence on the larval caste fate

To determine whether trophic eggs influence the process of caste determination, we compared the development of freshly hatched (first instar) larvae placed in small recipient colonies with and without trophic eggs. From each of 22 donor colonies, we obtained approximately 30 freshly hatched larvae by isolating the queens for 16 hours (from 2pm to 6am) every day for three weeks (in October 2020), with a 24-hour break every three days. Eggs were collected every eight hours and placed during 10 days in a petri-dish with a water reservoir ensuring a high humidity until they hatched. After hatching, for each colony, half of the larvae (i.e., 15 larvae) were then transferred into a recipient colony containing 20 workers, while the other half were placed in identical recipient colonies, which received in addition 45 0-4 hours-old trophic eggs (i.e., there were 3 trophic eggs per larva). There was no cross-fostering between colonies, so that larvae were always placed in recipient colonies containing workers from the same donor colony. The recipient colonies were maintained at 28°C and 60% humidity, with a 12-h/12-h light:dark cycle and fed *ad libitum* twice a week with grass seeds, flies and 20% honey water. The caste of each newly-produced individual was recorded at the pupal stage. To compare the proportion of queen pupae produced between recipient colonies with and without trophic eggs, we used the package *lme4* (Bates *et al*. 2015) to perform a binomial generalized linear mixed effects analysis (GLMM/link=“logit”) fit by maximum likelihood, with caste as response variable (binary categorical factor) and presence/absence of trophic eggs as an explanatory variable. Donor colony was included as a random effect. To test whether the presence of trophic eggs affects survival, we performed a linear mixed effect analysis with mortality as a response variable, presence/absence of trophic eggs as explanatory variable, and colonies as random effects. As we found a significantly higher survival of larvae in recipient colonies with trophic eggs than recipient colonies without trophic eggs (see results), we tested whether the percentage of larvae developing into queens was correlated with survival by performing a linear mixed effects analysis with the percentage of queen pupae as response variable, the survival as an explanatory variable and colonies as a random factor.

### Volume and content of trophic and viable eggs

The volumes of trophic (n=11) and viable eggs (n=14) were estimated by using the volume of an ellipse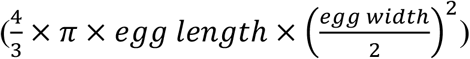, with egg length and width estimated on images under 10x magnification using ZEN Microscopy Software (v. 1.1.2.0).

To determine the nutritional content of viable and trophic eggs, we quantified the proteins, triglycerides, glycogen, and glucose in both types of eggs. We also quantified long and small RNAs (including miRNAs) as these compounds have been shown to be involved in caste determination in other eusocial species. To obtain the two types of eggs, we isolated 12 queens for 10 hours (7am to 5pm; from March to October 2021) in a dark petri-dish with three workers and a water supply. Eggs were collected every hour (so all eggs were a maximum of one hour old), and trophic and viable eggs were flash-frozen separately in liquid nitrogen. Twenty eggs were pooled for triglycerides-sugar-protein analyses and six eggs for RNA analyses. They were kept at -80°C until the extractions were performed. After the 10 hours of isolation, queens and workers were returned to their colony until the next isolation session. For each of the 12 colonies, we obtained two replicates of viable and trophic egg pools (i.e., 24 replicates in total).

Triglycerides, glycogen and glucose were quantified as described in Tennessen *et al*. (2014), and protein levels were measured using a Bradford assay (Bradford 1976). The 20 one-hour old eggs per sample were homogenized with beads in 200μl of PBS buffer in a Precellys Evolution tissue homogenizer coupled with a Cryolys Evolution (Bertin Technologies SAS).

For the Bradford assay, 10µl of the homogenate were put in a clear-bottom 96-well plate with 300µl of Coomassie Plus Reagent (Thermo Scientific: 23200) and incubated for 10 minutes at room temperature. Protein standard (Sigma: P5369) was used as standard (ranging from 0-0.5mg/ml) and protein absorbance was read at 595nm on a Hidex Sense Microplate Reader.

For the triglycerides assay, 90µl of homogenate were heat treated at 70°C for 10 minutes, then 40µl were mixed with 40µl of Triglyceride Reagent (Sigma: T2449) for digestion and 40µl were mixed with PBS buffer for free glycerol measurement. After 30 minutes incubation at 37°C, 30µl of each sample and standards were transferred to clear-bottom 96-well plate. 100µl of Free Glycerol Reagent (Sigma: F6428) was added to each sample, mixed well by pipetting, and incubated five minutes at 37°C. Glycerol standard solution (Sigma: G7793) was used as standard (ranging from 0-1.0mg/ml TAG) and absorbance was read at 540nm on a Hidex Sense Microplate Reader. The triglycerides concentration in each sample was determined by subtracting the absorbance of free glycerol in the corresponding sample.

Glucose and glycogen were quantified as in Tennessen *et al*. (2014). A 90µl aliquot was heat treated at 70°C for 10min and then diluted 1:2 with PBS. The standard curves for glucose (Sigma, G6918) and glycogen (Sigma: G0885) were made by diluting stocks to 160µg/ml, making 1:1 serial dilution for 160, 80, 40, 20 and 10µg/ml. 40µl of each sample was pipetted in duplicates of a clear microplate, and 30µl of each glucose or glycogen standard was pipetted in duplicates. Amyloglucosidase enzyme (Sigma, A1602) was diluted 3µl into 2000µl of PBS, and 40µl diluted enzyme was pipetted to the glycogen standards and to one well of the sample (for total glucose determination), 40µl PBS was pipetted to the glucose standards and to the other sample well (for free glucose determination). The plate was incubated at 37°C for 60 minutes. 30µl of each standard and samples (in duplicates) were transferred to a UV 96-well plate and 100µl Glucose Assay Reagent (G3293) was pipetted to each well. The plate was incubated at room temperature for 15 minutes and the absorbance was read at 340nm on a Hidex Sense Microplate Reader. The glycogen concentration was quantified by subtracting the free glucose absorbance from the total glycogen + glucose absorbance.

Concentrations of each compound (protein, triglycerides, glycogen, and glucose) were compared between viable and trophic eggs using a linear mixed effects analysis (LMER; package *lme4*), with the concentration as response variable and egg type as explanatory variable. Colony and extraction batch were added as random effects in the model.

### Total and small RNA, and DNA

RNA (>200 nt) and small RNA were isolated using the miRNeasy Mini Kit (Qiagen, cat. no. 217004) and RNeasy® MinElute® Cleanup Kit (Qiagen, cat. no. 74204), respectively, following manufacturer instructions. RNA (>200 nt) and small RNA concentrations were measured with a QuantiFluor® RNA System (Promega). RNA (>200 nt) integrity was examined with an Agilent Fragment Analyzer (at the Lausanne Genomic Technologies Facility) using a High Sensitivity Assay and small RNA were examined using the small RNA kit (at the Gene Expression Core Facility at EPFL).

The miRNA and RNA (>200 nt) concentrations were compared between viable and trophic eggs with paired-t-tests (for each type of eggs we used the average of the two replicates per colony). We also compared the fragment size distributions from 18 to 24 nucleotides for miRNAs (Sohel 2016) with a PCA and a Mantel test.

DNA was extracted from pools of six eggs using TRIzol (Life Technologies). DNA concentration was measured with a Nanodrop 3300 (ThermoFisher), and DNA integrity was examined with an Agilent Fragment Analyzer (at the Lausanne Genomic Technologies Facility) using a High Sensitivity Assay. DNA concentrations were compared between viable and trophic eggs using paired-t-tests (sample size is 5 for both types of eggs, each sample being a pool of 6 eggs).

## Results

### Trophic and viable egg characteristics

*P. rugosus* queens lay two types of eggs that are morphologically different. Viable eggs are white with a bright surface and have a distinct oval shape, a homogenous content as well as a solid chorion (Figure 2A), while trophic eggs are rounder, have a smooth surface and a granular looking content as well as a fragile chorion (Figure 2D). Trophic eggs had a significantly larger volume (94.3±4.3nL; n=11) than viable eggs (n=14; 63.3±1.6nL; two-sample t-test, t(23) = -9.54, p = 1.8×10^−09^). While viable eggs showed embryonic development at 25 and 65 hours (Fig 12B, C) there was no such development for trophic eggs (Fig. 2E, F). *P. rugosus* workers only laid viable eggs. They started to lay eggs approximately three weeks after queen removal (n=12 queenless recipient colonies) and approximately 90% of the eggs successfully hatched. However, only approximately 5% successfully developed into pupae which were all males.

**Figure 2.**
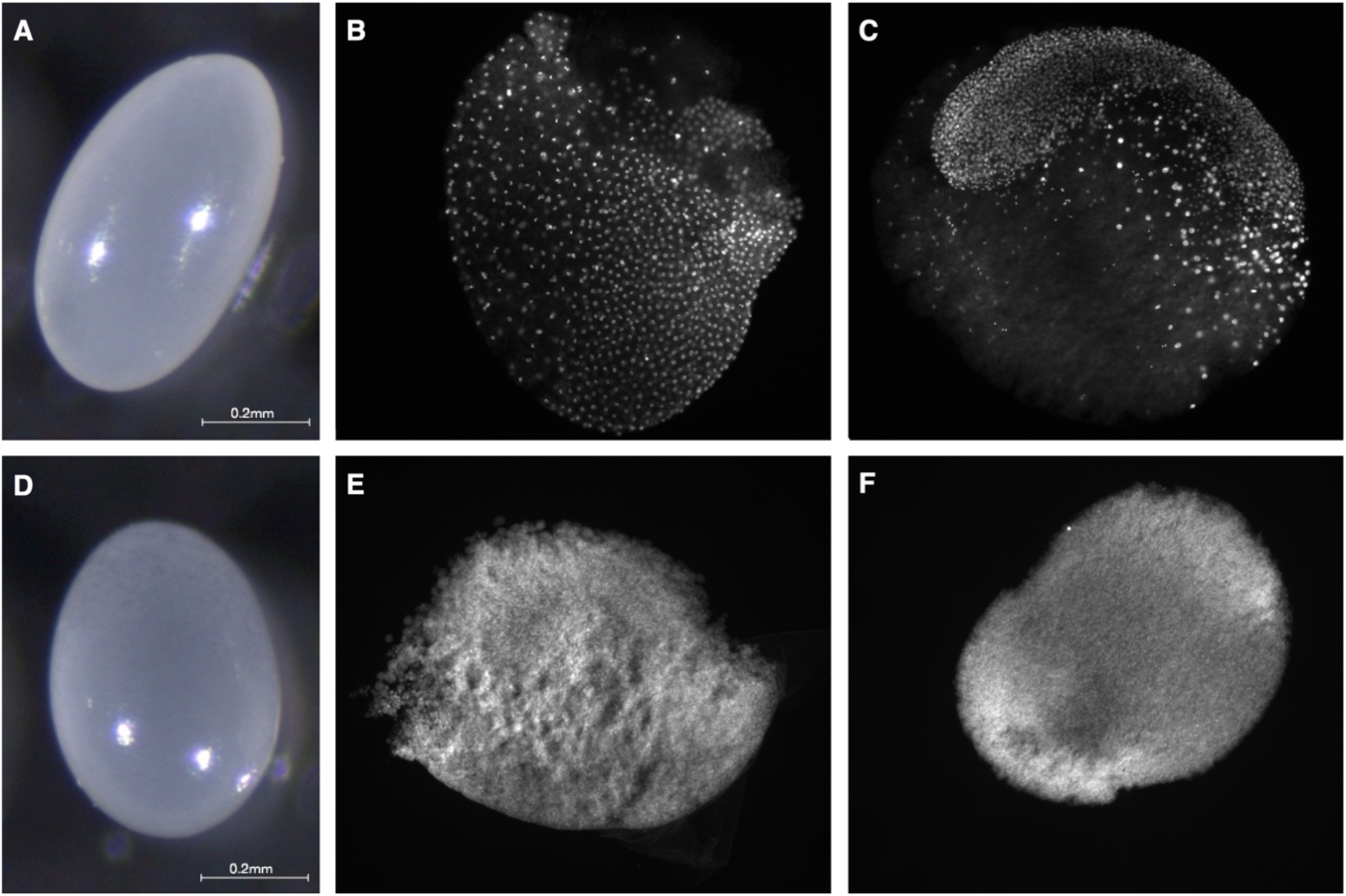
Morphology and of viable (**A**) and trophic (**D**) eggs laid by *P. rugosus* queens. Fluorescence images with DAPI-counterstained nuclei showing embryonic development of viable eggs at approximately 25 hours (**B**) and 65 hours (**C**). For trophic eggs there was no embryonic development at 25 hours (**E**) nor at 65 hours (**F**).

The percentage of eggs that were trophic was higher before hibernation (61.6±1.4% mean± SE; n=43 colonies) than after (50.3±2.0%; LMER, t(86)=5.04, p=9×10^−6^). The production of the two types of eggs was not random (Wald-Wolfowitz runs tests, p-values for the 11 queens in Table 1). Instead, each of the 11 queens tended to lay relatively long sequences of either viable (6.1±0.7; mean number per sequence ±SE) or trophic eggs (6.0±0.5; Figure 3).

**Table 1.** Wald-Wolfowitz runs tests on the queen’s egg sequence. Significant p-values (corrected for multiple testing) indicate that queens do not lay viable and trophic eggs in a random sequence.

| Queen ID | P-value for egg sequence | P-value of random sequence | Number of eggs per sequence |
| --- | --- | --- | --- |
| **338** | $4.1 \times 10^{-11}$ | 0.419 | 94 |
| **117** | $4.3 \times 10^{-07}$ | 0.567 | 63 |
| **173** | $1.7 \times 10^{-03}$ | 0.755 | 92 |
| **303** | $4.8 \times 10^{-13}$ | 0.765 | 110 |
| **215** | $9.8 \times 10^{-11}$ | 0.292 | 70 |
| **120** | $1.4 \times 10^{-05}$ | 0.518 | 75 |
| **12B** | $1.9 \times 10^{-03}$ | 0.298 | 38 |
| **316** | $3.4 \times 10^{-09}$ | 0.737 | 93 |
| **193** | $4.3 \times 10^{-12}$ | 0.655 | 101 |
| **150** | $1.4 \times 10^{-10}$ | 0.630 | 62 |
| **125** | $1.5 \times 10^{-05}$ | 0.404 | 58 |

**Figure 3.**
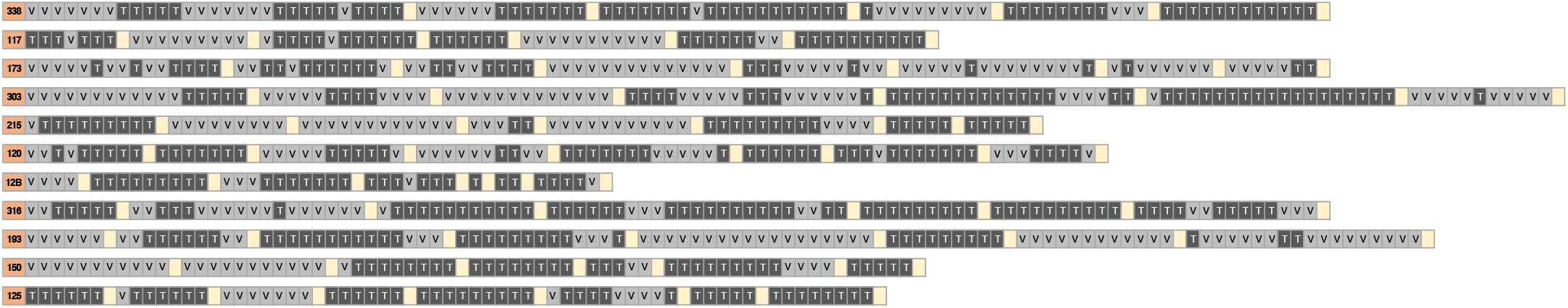
Egg-laying sequences from eleven *P. rugosus* queens. Every row shows the sequence of viable (V) and trophic (T) eggs laid by a given queen (queen ID in the orange cell). Each egg laying session lasted 10 hours. The yellow squares indicate the intervals (16 hours to several days) between egg-laying sessions.

The concentrations of protein, triglycerides, glycogen, and glucose were significantly higher in viable than trophic eggs (LMER, protein: t = -13.11, p <0.0001; triglycerides: t = -11.66, p <0.0001; glycogen: t = -11.98, p <0.0001; glucose: t = -18.60, p <0.0001; Figure 4).

**Figure 4.**
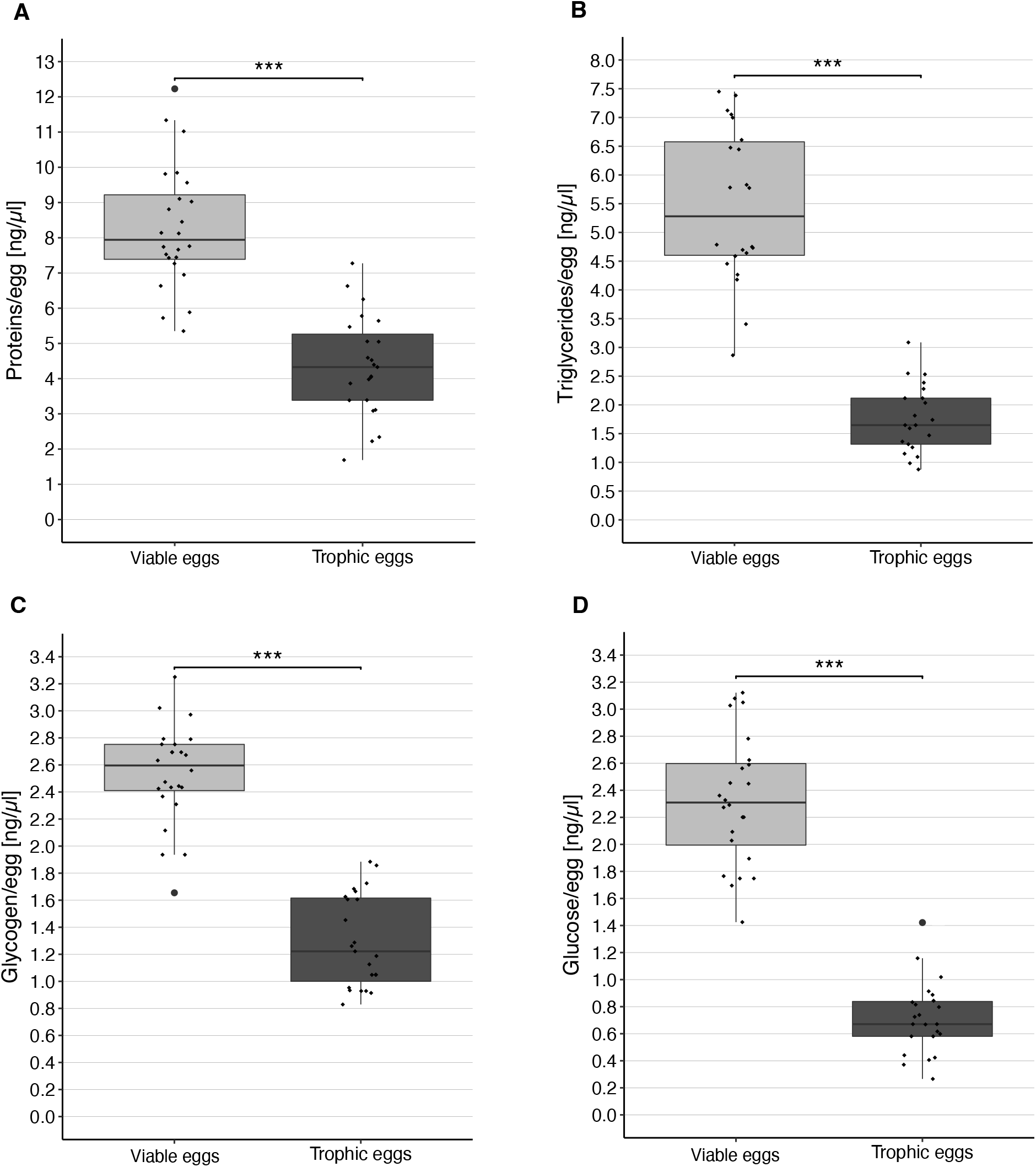
Concentration (± standard error) of protein (**A**), triglycerides (**B**), glycogen (**C**) and glucose (**D**) in viable and trophic eggs. Each dot represents the average of the two replicates per colony.

The amount of small RNA (<200 nt, including miRNA and tRNA; Nagano and Fraser 2011) was significantly higher in viable eggs (44.3±1.4ng, mean ± SE) than in trophic eggs (22.3±1.1ng; paired-t-test, t_(23)_ = 15.9, p = 6.5*10^−14^). The same was true for longer RNAs (>200 nt; viable eggs: 7.6±0.6ng, mean ± SE; trophic eggs: 3.6±0.3ng; paired-t-test, t_(23)_ = 7.2, p = 2.7*10^−7^).

The DNA quantification showed that the amount of DNA was about twice higher in viable (15.9±1.9ng/µl) than trophic eggs (8.8±1.9ng/µl; t-test, t_(4.7)_ = 2.7, p = 0.045).

There was a significant difference in the miRNA fragment size distribution between viable and trophic eggs (Mantel test, r_M_ = 0.26, p<.0001), as shown on the PCA (Figure 5A). There was no difference in the tRNA fragment size distribution between the two types of eggs (Mantel test, r_M_ = 0.01, p=0.30, Figure 5B).

**Figure 5.**
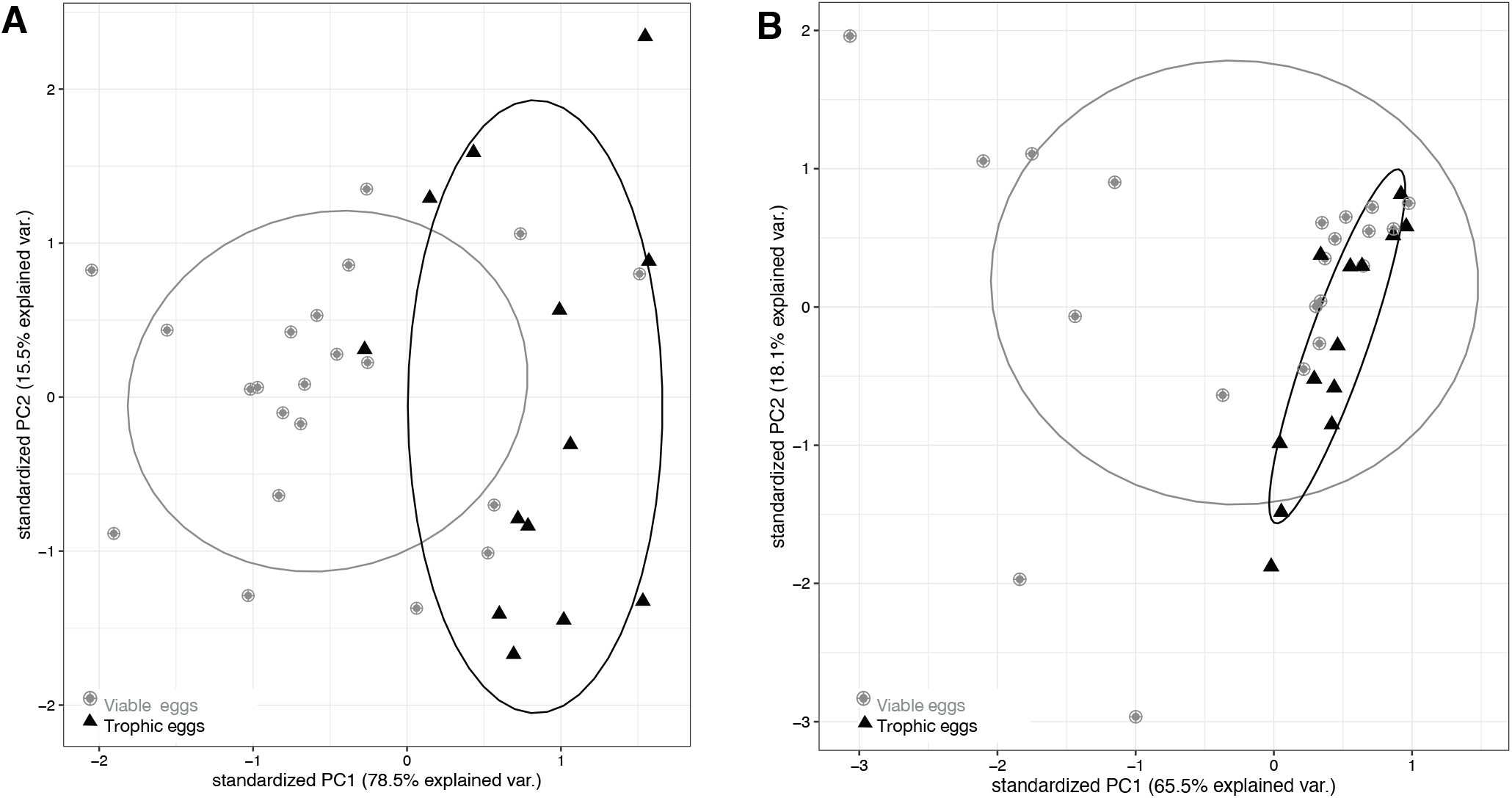
First two principal components (PC1 and PC2) explaining size distribution variation for (**A**) miRNA and (**B**) tRNA across egg samples, with viable eggs in grey dots and trophic eggs in black triangles. Ellipses enclose each of the egg type groups.

### Trophic eggs influence caste fate of larvae

The percentage of larvae that developed into queens was significantly lower in recipient colonies that received trophic eggs (27±9% mean±SE; n=22) than in recipient colonies without trophic eggs (83±10%; n=22; binomial GLMM (link=“logit”), z = 4.25, p = 2×10^−5^; Figure 6A and Supplementary Table 2). The survival of larvae until the pupal stage was also significantly lower in the colonies without trophic eggs (16.9±3.8%; n=22; LMER, z = 2.66, p = 0.008) than in colonies with trophic eggs (30.2±6.7%; mean±SE; n=22), but the 1.8 fold survival decrease cannot fully account for the 3 fold difference in queen percentage between the two treatments. Furthermore, there was no significant correlation between larval mortality and the percentage of larvae developing into queens (n=44 recipient colonies; LMER, z = 0.97, p = 0.34; Figure 6B). These analyses allow us to exclude differential survival between castes as the sole explanation for the higher percentage of queens developing in the recipient colonies without trophic eggs.

**Figure 6.**
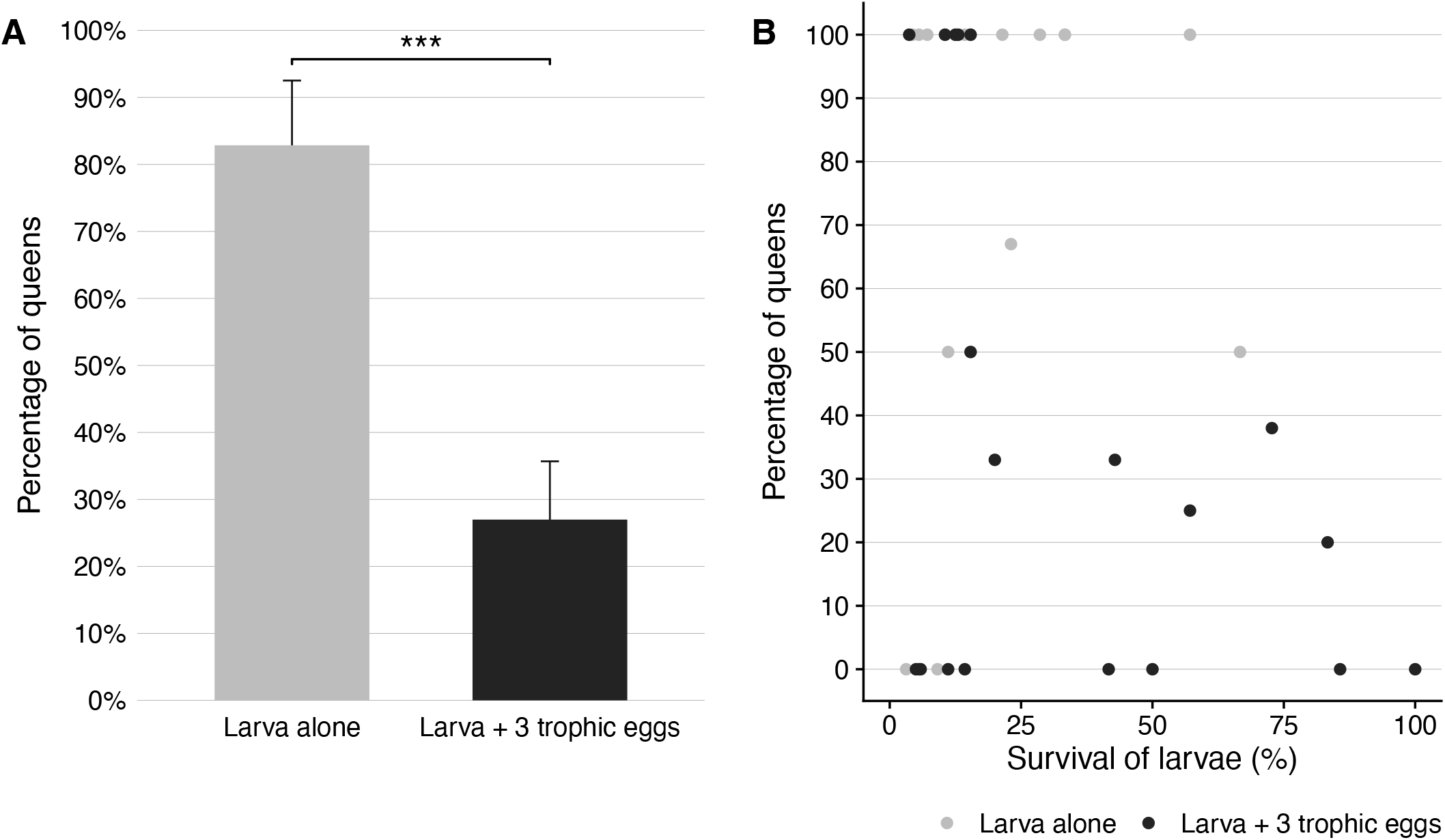
(**A**) Percentage (± standard error) of queens among the larvae that developed to the pupal stage in colonies without (grey) or with (black) trophic eggs. (**B**) Relationship between the percentage of larvae which developed into queens and the survival of larvae (percentage) between the larval to pupal stages.

## Discussion

Our study reveals that *P. rugosus* queens lay a very high proportion (0.6) of trophic eggs. These eggs differ in many ways from viable eggs. First, trophic eggs are larger, rounder, have a smoother surface, a more granular looking content as well as a more fragile chorion than viable eggs. Similar differences between trophic and viable eggs have been reported in other ant species (Wilson 1976; Wardlaw and Elmes 1995; Gobin *et al*. 1998; Dietemann and Peeters 2000; Dietemann *et al*. 2002; Perry and Roitberg 2006; Lee *et al*. 2017). Our analyses also showed that trophic eggs are solely laid by queens; *P. rugosus* workers are able to produce viable eggs which occasionally develop into males, but they do not lay trophic eggs. Moreover, trophic eggs have a reduced DNA content.

Importantly, our experiments showed that the presence of trophic eggs influences the process of caste determination. First instar female larvae fed with trophic eggs were significantly more likely to develop into workers than larvae without access to trophic eggs. This was somewhat surprising because trophic eggs are generally thought to be an important source of nutrients to the colony and, everything else being equal, one would think that eating such eggs should increase the likelihood of females to develop into queens (which are usually larger than workers). Indeed, two earlier studies suggested that an increase in the proportion of trophic eggs might be associated with an increase in the proportion of larvae developing into queens (*L. humile*: Bartels 1988; *P. barbatus*: Helms Cahan *et al*. 2011). However, in these two studies variation in the availability of trophic eggs was associated with other differences (number of queens in the colony, Bartels 1988; Administration of a JH analogue, Helms Cahan *et al*. 2011) making it difficult to determine what factor had a causal effect. It would be interesting to experimentally manipulate the quantity of trophic eggs in *L. humile* and *P. barbatus* to determine whether they have a positive or inhibitory effect on the likelihood of larvae to develop into queens.

Our analyses revealed that trophic eggs have a lower content of protein, triglycerides, glycogen, and glucose than viable eggs. A reduced protein content of trophic as compared to viable eggs has also been documented in *Pheidole pallidula* (Lorber and Passera 1981). These findings are in line with the view that trophic eggs do not simply have a nutritional function as it might then be expected that they should at least contain as much nutrients as viable eggs. Interestingly, our analyses also revealed important differences in RNA and miRNA content between the two egg types. miRNAs have already been suggested to influence larval caste determination in the honeybee (Guo *et al*. 2013) with worker jelly being enriched in miRNAs compared to royal jelly (Guo *et al*. 2013; Zhu *et al*. 2017). These studies suggest that it is not the royal jelly that stimulates larval determination into queen, but rather the worker jelly which stimulates the development of larvae into workers. Similarly, our study reveals that compounds found in trophic eggs, perhaps miRNAs, influence larval development towards the worker phenotype. Interestingly, it has also been recently shown that trophallactic fluid in the ant *Camponotus floridanus* contains non-digestive related proteins, microRNAs and juvenile hormone (LeBoeuf *et al*. 2016). Moreover, comparison of trophallactic fluid proteins across social insect species revealed that many are regulators of growth, development and behavioral maturation (Meurville and LeBoeuf 2021). Finally, a recent study showed that pupae of several ant species produce secretions that play an important role for early larval nutrition with young larvae exhibiting stunted growth and decreased survival without access to the fluid (Snir *et al*. 2022). This raises the possibility that chemicals delivered in trophic eggs, trophallactic fluids and pupae secretions play previously unsuspected roles in communication and caste development. Given that some ants do not perform trophallaxis, it would be interesting to determine whether there are differences in the content of trophic eggs of species performing trophallaxis and species which do not.

Maternal effects on the process of caste determination have been demonstrated in several social insect species, including *P. rugosus*, either by queen behaviour or content of the eggs being produced (De Menten *et al*. 2005; Linksvayer 2006; Schwander *et al*. 2008b; Libbrecht *et al*. 2013; Wei *et al*. 2019). This is, to our knowledge the first experimental demonstrations that provisioning of trophic eggs influences caste fate. Since only queens produce trophic eggs in *P. rugosus*, trophic egg provisioning could be the main mechanism underlying the previously documented maternal effects on the process of caste determination. In species where workers produce trophic eggs (Supplementary Table 1), the same mechanism could allow workers to influence colony level caste ratios if their eggs do also affect the process of caste determination. The presence of trophic eggs has been documented in only relatively few ant species (Table 1). However, their presence has been investigated in only few species and it is likely that trophic eggs are produced in most ant species, particularly in those with an independent mode of colony founding,

Finally, our analyses also revealed seasonal differences in the proportion of viable and trophic eggs, with a higher ratio of trophic eggs before hibernation than after. In *Pogonomyrmex* the production of new queens occurs after hibernation (Smith and Tschinkel 2006) or when the queen dies or is removed from the colony (pers. obs). Thus, new queens are typically produced when there are fewer trophic eggs. Our results predict that under natural conditions, a decrease in the proportion of trophic eggs should lead to an increase in the proportion of larvae developing into queens. The same logic applies to species where trophic eggs are laid only by the workers in queenright colonies (Supplementary Table 1). After the queen’s death, workers start producing their own male offspring and lay mostly (if not only) viable eggs (*Temnothorax recedens*, Dejean and Passera 1974; *Plagiolepis pygmaea*, Passera 1980; *Myrmecia gulosa*, Dietemann *et al*. 2002), which again leads to a decrease, or cessation, in trophic egg production. A decrease in trophic egg production and the development of queens were observed simultaneously in freshly orphaned colonies of *Temnothorax recedens* (Dejean and Passera 1974), *Plagiolepis pygmaea* (Passera 1980) and *Myrmecia gulosa* (Dietemann *et al*. 2002). These examples are consistent with the view that trophic eggs may also play a role in the process of caste determination in other ant species.

In addition to the seasonal differences reported in this study, previous studies in ants have also revealed that young queens produce more trophic eggs than older queens (Hölldobler & Wilson 1990). Interestingly, the production of new queens in ant colonies start only when colonies are several years old and have reached a relatively large size. Thus, this raises the possibility that the decrease of the proportion of trophic eggs laid by queens when they age may contribute to the production of new queens being restricted to colonies having reached a relatively large size.

Egg cannibalism has been reported in a many ant species (Wilson 1971; Sorensen *et al*. 1983; Bourke 1991; Crespi 1991; Peeters and Tsuji 1993; Aron *et al*. 1994; Heinze *et al*.1999; Schultner *et al*. 2013) and may serve many purposes, including preferential elimination of males, source of energy (Sorensen *et al*. 1983) and intracolony conflict (Schultner *et al*. 2014). Our study indicates a new potential role of egg cannibalism as a mechanism to regulate the allocation of energy into the production of new queens versus workers.

In conclusion, this study provides a new striking example of how females can influence the developmental fate of their offspring. Because many ants produce trophic eggs, it is possible that this mechanism of parental manipulation is widespread and plays an important role in the general process of caste determination. It would be interesting to conduct manipulative experiments similar to those of this study in additional species, to determine whether trophic eggs broadly play a role in the process of caste determination. Of interest would also be to determine what chemicals in the egg are responsible for influencing the development of larvae.

## Acknowledgments

We thank Dr. C. Berney for technical assistance to develop wet lab protocols. We are grateful to S. McGregor and M. Chapuisat for their helpful comments on the manuscript. This work was supported by an ERC grant and the Swiss NSF (LK) and funding from the University of Lausanne (LK and TS).

## References

Amor, F., P. Ortega, R. Boulay, and X. Cerdá. 2017. Frequent colony orphaning triggers the production of replacement queens via worker thelytoky in a desert-dwelling ant. Insectes Soc. 64:373–378. doi.org/10.1007/s00040-017-0556-9

Aron, S., Passera, L. Keller, L. 1994. Queen-worker conflict over sex ratio: A comparison of primary and secondary sex ratios in the Argentine ant, Iridomyrmex humilis. J. Evol. Biol. 7:403–418

Augustin, J. O., J. F. Santos, and S. L. Elliot. 2011. A behavioral repertoire of Atta sexdens (Hymenoptera, Formicidae) queens during the claustral founding and ergonomic stages. Insectes Soc. 58:197–206. doi.org/10.1007/s00040-010-0137-7

Azevedo, D. O., J. C. Zanuncio, J. H. C. Delabie, and J.E. Serrão. 2011. Temporal variation of vitellogenin synthesis in Ectatomma tuberculatum (Formicidae: Ectatomminae) workers. J. Insect Physiol. 57:972–977. doi.org/10.1016/j.jinsphys.2011.04.015

Baroni Urbani, C. B. 1991. Indiscriminate oophagy by ant larvae: an explanation for brood serial organization? Insectes Soc. 38:229–239. doi.org/10.1007/BF01314909

Bartels, P. J. 1988. Reproductive caste inhibition by Argentine ant queens: New mechanisms of queen control. Insectes Soc. 35:70–81. doi.org/10.1007/BF02224139

Bates, D., M. Maechler, B. Bolker, and S. Walker. 2015. Fitting Linear Mixed-Effects Models Using lme4. J. Stat. Softw. 67:1–48. doi.org/doi:10.18637/jss.v067.i01

Blake, J. A., and P. L. Arnofsky. 1999. Reproduction and larval development of the spioniform Polychaeta with application to systematics and phylogeny. Hydrobiologia. 402:57–106. doi.org/10.1023/A

Bourke, A. F. G. 1988. Worker reproduction in the higher eusocial Hymenoptera. Quart Rev Biol 63: 291–311.

Bourke, A. F. G. 1991. Queen behaviour, reproduction and egg cannibalism in multiple-queen colonies of the ant Leptothorax acervorum. Animal Behav. 42:295–310.

Bradford, M. M. 1976. A rapid and sensitive method for the quantification of microgram quantities of protein utilizing the principle of protein–dye binding. Anal. Biochem. 72:248–254. doi.org/10.1016/0003-2697(76)90527-3

Brian, M. V, Rigby, C. 1978. The trophic eggs of Myrmica rubra L. Insect Molecular Biology. 25(1):89–110.

CamargoMathias, M., and F. Caetano. 1995. Trophic eggs in workers of Neoponera villosa ants (Hymenoptera:Ponerinae). J. Adv. Zool. 16:62–66.

Cameron, R. C., E. J. Duncan, and P. K. Dearden. 2013. Biased gene expression in early honeybee larval development. BMC Genomics. 14:1–12. BioMed Central. doi.org/10.1186/1471-2164-14-903

Cassill, D. 2002. Brood care strategies by newly mated monogyne Solenopsis invicta (Hymenoptera: Formicidae) queens during colony founding. Ann. Entomol. Soc. Am. 95:208–212. doi.org/10.1603/0013-8746(2002)095[0208:BCSBNM]2.0.CO;2

Choe, J. C. 1988. Worker reproduction and social evolution in ants (Hymenoptera: Formicidae). Advances in myrmecology. 163–187

Collin, R. 2004. Phylogenetic effects, the loss of complex characters, and the evolution of development in calyptraeid gastropods. Evolution. 58:1488–1502. doi.org/10.1111/j.0014-3820.2004.tb01729.x

Collins, D. H., I. Mohorianu, M. Beckers, V. Moulton, T. Dalmay, and Bourke A. F. 2017. MicroRNAs Associated with Caste Determination and Determination in a Primitively Eusocial Insect. Sci. Rep. 7:1–9. Nature Publishing Group. doi.org/10.1038/srep45674

Crespi, B. J. 1977. Cannibalism and trophic eggs in subsocial and eusocial insects. Pp. 176–213 in M. A. Elgar and B. J. Crespi, eds. Cannibalism: Ecology and Evolution Among Diverse Taxa.

Dejean, A., and L. Passera. 1974. Ponte des ouvrières et inhibition royale chez la Fourmi Temnothorax recedens (Nyl.) (Formicidae, Myrmicinae). Insectes Soc. 21:343–355. doi.org/10.1007/BF02331564

Della Lucia, T. M. C., E. F. Vilela, D. D. O. Moreira, J. M. S. Bento, and N. Dos Anjos. 1990. Egg-laying in Atta sexdens rubropilosa, under laboratory conditions. Pp. 173–179 in R. K. Vander Meer, K. Jaffé, and A. Cedeno, eds. Applied Myrmecology: A World Perspective. Westview Press, New York.

Dietemann, V., B. Hölldobler, and C. Peeters. 2002. Caste specialization and determination in reproductive potential in the phylogenetically primitive ant Myrmecia gulosa. Insectes Soc. 49:289–298. doi.org/10.1007/s00040-002-8316-9

Dietemann, V., and C. Peeters. 2000. Queen influence on the shift from trophic to reproductive eggs laid by workers of the ponerine ant Pachycondyla apicalis. Insectes Soc. 47:223–228. doi.org/10.1007/PL00001707

Dijkstra, M. B., D. R. Nash, and J. J. Boomsma. 2005. Self-restraint and sterility in workers of Acromyrmex and Atta leafcutter ants. Insectes Soc. 52:67–76. doi.org/10.1007/s00040-004-0775-8

Fletcher, D. J. C., and K. G. Ross. 1985. Regulation of reproduction in Eusocial hymenoptera. Annu. Rev. Entomol. 30:319–343.

Freeland, J. 1958. Biological and social patterns in the Australian bulldog ants of the genus Myrmecia. Aust. J. Zool. 6:1–18. doi.org/10.1071/ZO9580001

Gibson, G., C. Hart, C. Coulter, and H. Xu. 2012. Nurse eggs form through an active process of apoptosis in the spionid Polydora cornuta (Annelida). Integr. Comp. Biol. 52:151–160. doi.org/10.1093/icb/ics054

Gobin, B., J. Billen, and C. Peeters. 1999. Policing behaviour towards virgin egg layers in a polygynous ponerine ant. Anim. Behav. 58:1117–1122. doi.org/10.1006/anbe.1999.1245

Gobin, B., and F. Ito. 2000. Queens and major workers of Acanthomyrmex ferox redistribute nutrients with trophic eggs. Naturwissenschaften 87:323–326. doi.org/10.1007/s001140050731

Gobin, B., C. Peeters, and J. Billen. 1998. Production of trophic eggs by virgin workers in the ponerine ant Gnamptogenys menadensis. Physiol. Entomol. 23:329–336. doi.org/10.1046/j.1365-3032.1998.00102.x

Guo, X., S. Su, G. Skogerboe, S. Dai, W. Li, Z. Li, F. Liu, R. Ni, Y. Guo, S. Chen, S. Zhang, and R. Chen. 2013. Recipe for a busy bee: MicroRNAs in honey bee caste determination. PLoS One 8:1– 10. doi.org/10.1371/journal.pone.0081661

Hammond, R and L. Keller 2004. Conflict over male parentage in social insects. PLoS Biology, 2(9): e248. 1472-1482.

Haskins, C.P., and Whelden R.M. 1965. “Queenlessness,” Worker Sibship, and Colony Versus Population Structure in the Formicid Genus Rhytidoponera’. Psyche: A Journal of Entomology. 72(ID 040465): 26 pages. doi.org/10.1155/1965/40465

Heinze, J., S. Cover, and B. Hölldobler. 1995. Neither worker, nor queen: an ant caste specialized in the production of unfertilized eggs. Psyche A J. Entomol. 102:173–185. doi.org/10.1155/1995/65249

Heinze, J., S. Foitzik, B. Oberstadt, O. Rüppell, and B. Hölldobler. 1999. A female caste specialized for the production of unfertilized eggs in the ant Crematogaster smithi. Naturwissenschaften. 86:93– 95. doi.org/10.1007/s001140050579

Helms Cahan, S., C. J. Graves, and C. S. Brent. 2011. Intergenerational effect of juvenile hormone on offspring in Pogonomyrmex harvester ants. J. Comp. Physiol. B. 181:991–999. doi.org/10.1007/s00360-011-05Helms

Helms Cahan, S. and L. Keller L. 2003 Complex hybrid origin of genetic caste determination in harvester ants. Nature. 424:306–309

Helms Cahan S., J.D. Parker, S.W. Rissing, R.A. Johnson, T.S. Polony, M.D. Weiser and D. R. Smith D.R. 2002. Extreme genetic differences between queens and workers in hybridizing Pogonomyrmex harvester ants. Proc. R. Soc. Lond. B. Biol. Sci. 269:1871–1877

Hölldobler, B., and N. F. Carlin. 1989. Colony founding, queen control and worker reproduction in the ant Aphaenogaster (=Novomessor) cockerelli (Hymenoptera: Formicidae). Psyche (Stuttg). 96:131–152.

Hölldobler, B., and E. O. Wilson. 1983. Queen Control in Colonies of Weaver Ants (Hymenoptera: Formicidae). Ann. Entomol. Soc. Am. 76:235–238. doi.org/10.1093/aesa/76.2.235

Hölldobler, B., and E. O. Wilson. 1990 The Ants. Cambridge, MA: Belknap Press of Harvard Univ. Press.

Hora, R. R., C. Poteaux, C. Doums, D. Fresneau, and R. Fénéron. 2007. Egg cannibalism in a facultative polygynous ant: Conflict for reproduction or strategy to survive? Ethology. 113:909–916. doi.org/10.1111/j.1439-0310.2007.01391.x

Huber, J. 1905. Uber die Koloniegründung bei Atta sexdens. Biologisches Centralblatt 25:625–635.

Hung, A. C. F. 1973. Reproductive biology in dulotic ants: preliminary report (Hymenoptera: Formicidae). Entomol. News. 84:253–259.

Ito, F. 1991. Preliminary report on queenless reproduction in a primitive ponerine ant Amblyopone sp. (reclinata group) in west Java, indonesia. Psyche (Stuttg). 98:319–322.

Ito, F. 1993. Social organization in a primitive ponerine ant: Queenless reproduction, dominance hierarchy and functional polygyny in Amblyopone sp. (reclinata group) (hymenoptera: Formicidae: Ponerinae). J. Nat. Hist. 27:1315–1324. doi.org/10.1080/00222939300770751

Ito, F. 2005. Mechanisms regulating functional monogyny in a Japanese population of Leptothorax acervorum (Hymenoptera, Formicidae): Dominance hierarchy and preferential egg cannibalism. Belgian J. Zool. 135:3–8.

Julian, G.E. J.H., Fewell, J., Gadau J. R.A.,, Johnson R.A. and D. Larrabee D. 2002. Genetic determination of the queen caste in an ant hybrid zone. Proc. Natl. Acad. Sci. USA. 99:8157–8160

Keller, L. and L. Passera. 1989. Size and fat content of gynes in relation with the mode of colony founding in ants (Hymenoptera; Formicidae). Oecologia 80:236–240

Khila, A., and E. Abouheif. 2008. Reproductive constraint is a developmental mechanism that maintains social harmony in advanced ant societies. Proc. Natl. Acad. Sci. U.S.A. 105:17884–17889. doi.org/10.1073/pnas.0807351105

Khila, A., and E. Abouheif. 2010. Evaluating the role of reproductive constraints in ant social evolution. Philos. Trans. R. Soc. B. 365:617–630. doi.org/10.1098/rstb.2009.0257

Klowden, M. J. 2013. Developmental Systems. Pp. 149–196 in M. J. Klowden, ed. Physiological Systems in Insects (Third Edition). Academic Press.

Kudo, S., and T. Nakahira. 2004. Effects of Trophic-Eggs on Offspring Performance and Rivalry in a Sub-Social Bug. Oikos. 107:28–35.

LeBoeuf, A. C., P. Waridel, C. S. Brent, A. N. Gonçalves, L. Menin, D. Ortiz, O. Riba-Grognuz, A. Koto, Z. G. Soares, E. Privman, E. A. Miska, R. Benton, and L. Keller. 2016. Oral transfer of chemical cues, growth proteins and hormones in social insects. Elife. 5:e20375:1–27. doi.org/10.7554/eLife.20375

Lee, C. C., H. Nakao, S. P. Tseng, H. W. Hsu, G. L. Lin, J. W. Tay, J. Billen, F. Ito, C. Y. Lee, C. C. Lin, and C. C. S. Yang. 2017. Worker reproduction of the invasive yellow crazy ant Anoplolepis gracilipes. Front. Zool. 14(1):1–12. doi.org/10.1186/s12983-017-0210-4

Levin, L. A., and T. S. Bridges. 1995. Pattern and diversity in reproduction and development. Pp. 1–48 in L. McEdward, ed. Ecology of marine invertebrate larvae. CRC Press. doi.org/10.1201/9780138758950-1

Libbrecht, R., M. Corona, F. Wende, D. O. Azevedo, J. E. Serrão, and L. Keller. 2013. Interplay between insulin signaling, juvenile hormone, and vitellogenin regulates maternal effects on polyphenism in ants. Proc. Natl. Acad. Sci. U.S.A. 110:11050–11055. doi.org/10.1073/pnas.1221781110

Linksvayer, T. A. 2006. Direct, maternal, and sibsocial genetic effects on individual and colony traits in an ant. Evolution. 60(12):2552. doi.org/10.1554/06-011.1

López-Ortega, M., and T. Williams. 2018. Natural enemy defense, provisioning and oviposition site selection as maternal strategies to enhance offspring survival in a sub-social bug. PLoS One 13:1– 18. doi.org/10.1371/journal.pone.0195665

Lorber, B., and L. Passera. 1981. Étude comparative des protéines solubles des œufs de la fourmi Pheidole pallidula Nyl. Bull Intérieur SF-UIEIS 97–99.

Masuko, K. 2003. Larval oophagy in the ant Amblyopone silvestrii (Hymenoptera, Formicidae). Insectes Soc. 50:317–322. doi.org/10.1007/s00040-003-0688-y

De Menten, L., Fournier, D., Brent, C., Passera, L., Vargo, E.L., and Aron S. 2005. Dual Mechanism of Queen Influence over Sex Ratio in the Ant Pheidole Pallidula. Behavioral Ecology and Sociobiology. 58(6):527–33. doi.org/10.1007/s00265-005-0964-0.

Meurville, M.-P., and LeBoeuf, A.C. 2021. Trophallaxis: The Functions and Evolution of Social Fluid Exchange in Ant Colonies (Hymeno Ptera: Formicidae). Myrmecological News. 31:1–20. doi.org/doi:10.25849/myrmecol.news_031:00113

Nagano, T., and Fraser, P. 2011. No-Nonsense Functions for Long Noncoding RNAs. Cell 145:178–181. doi.org/10.1016/J.CELL.2011.03.014

Passera, L. 1978. Une nouvelle catégorie d’œufs alimentaires: les œufs alimentaires émis par les reines vierges de Pheidole pallidula (Nyl.) (Formicidae, Myrmicinae). Insectes Soc. 25:117–126.

Passera, L. 1980. La fonction inhibitrice des reines de la fourmi Plagiolepis pygmaea Latr.: Rôle des phéromones. Insectes Soc. 27:212–225. doi.org/10.1007/BF02223665

Peeters, C. 2017. Independent colony foundation in Paraponera clavata (hymenoptera: Formicidae): First workers lay trophic eggs to feed queen’s larvae. Sociobiology 64:417–422. doi.org/10.13102/sociobiology.v64i4.2092

Peeters, C. and Tsuji, K. 1993. Reproductive conflict among ant workers in Diacamma sp. from Japan: dominance and oviposition the absence of gamergate. Ins. Soc. 40:119–136

Perry, J. C., and B. D. Roitberg. 2006. Trophic egg laying: Hypotheses and tests. Oikos 112:706–714. doi.org/10.1111/j.0030-1299.2006.14498.x

Romiguier, J., Borowiec, M. L., Weyna, A., Helleu, Q., Loire, E., La Mendola, C., Rabeling, C., Fisher, B. L., Ward, P. S., Keller, L. 2022. Ant phylogenomics reveals a natural selection hotspot preceding the origin of complex eusociality. Current Biology 32(13):2942-2947.e4. doi.org/10.1016/j.cub.2022.05.001

RStudio Team. 2015. RStudio: Integrated Development Environment for R. Boston, MA.

Schultner, E., d’Ettorre, P., Helanterä, H. 2013, Social conflict in ant larvae: egg cannibalism occurs mainly in males and larvae prefer alien eggs, Behavioral Ecol., 24:1306–1311. doi.org/10.1093/beheco/art067.

Schultner, E., Gardner, A., Karhunen, M., Helanterä H. 2014. Ant larvae as players in social conflict: Relatedness and individual identity mediate cannibalism Intensity. Am. Nat. 184:E161–E174.

Schwander, T., Cahan, S. H. & Keller, L. 200). Characterization and distribution of Pogonomyrmex harvester ant lineages with genetic caste determination. Molecular Ecology 16:367–387.

Schwander, T., J. Y. Humbert, C. S. Brent, S. H. Cahan, L. Chapuis, E. Renai, and L. Keller. 2008. Maternal Effect on Female Caste Determination in a Social Insect. Curr. Biol. 18:265–269. doi.org/10.1016/j.cub.2008.01.024

Smeeton, L. 1981. The Source of Males in Myrmica Rubra L. (Hym. Formicidae). Insectes Sociaux. 28(3):263–78. doi.org/10.1007/BF02223628.

Smith, A. A., B. Hölldobler, and J. Liebig. 2008. Hydrocarbon signals explain the pattern of worker and egg policing in the ant Aphaenogaster cockerelli. J. Chem. Ecol. 34:1275–1282. doi.org/10.1007/s10886-008-9529-9

Smith, C. R., C. Schoenick, K. E. Anderson, J. Gadau, and A. V Suarez. 2007. Potential and realized reproduction by different worker castes in queen-less and queen-right colonies of Pogonomyrmex badius. Insect. Soc. 54:260–267. doi.org/10.1007/s00040-007-0940-y

Smith, C. R., & Suarez, A. V. (2010). The trophic ecology of castes in harvester ant colonies. Functional Ecology, 24(1), 122–130. doi.org/10.1111/j.1365-2435.2009.01604.x

Smith, C. R., and W. R. Tschinkel. 2006. The sociometry and sociogenesis of reproduction in the Florida harvester ant, Pogonomyrmex badius. J. Insect Sci. 6:1–11. doi.org/10.1673/2006_06_32.1

Snir, O., Alwaseem, H., Heissel, S. Sharma, A., Valdés-Rodríguez, S., Carroll, T.S., Jiang, C.S., Razzauti, J., Kronauer, D.J.C. 2022. The pupal moulting fluid has evolved social functions in ants. Nature. 612(7940):488–494. doi.org/10.1038/s41586-022-05480-9

Sohel, M. H. 2016. Extracellular/Circulating MicroRNAs: Release Mechanisms, Functions and Challenges. Achiev. Life Sci. 10:175–186. doi.org/10.1016/j.als.2016.11.007

Sorensen, A., T. Busch, and S. Vinson. 1983. Factors affecting brood cannibalism in laboratory colonies of the imported fire ant, Solenopsis invicta Buren (Hymenoptera: Formicidae). J. Kansas Entomol. Soc, 56:140–150

Søvik, E., G. Bloch, and Y. Ben-Shahar. 2015. Function and evolution of microRNAs in eusocial Hymenoptera. Front. Genet. 6:1–11. doi.org/10.3389/fgene.2015.00193

Strathmann, M. F., and R. R. Strathmann. 2006. A vermetid gastropod with complex intracapsular cannibalism of nurse eggs and sibling larvae and a high potential for invasion. Pacific Sci. 60:97– 108. doi.org/10.1353/psc.2005.0062

Suzzoni, J. P., L. Passera, and A. Strambi. 1979. Taux des ecdystéroides et déterminisme des castes de la Fourmi Pheidole pallidula (Hym. Formicidae).

Taylor, R.W. 1978. Nothomyrmecia Macrops : A Living-Fossil Ant Rediscovered. Science. 201(4360):979–85. doi.org/10.1126/science.201.4360.979

Tennessen, J. aso. M., W. E. Barry, J. Cox, and C. S. Thummel. 2014. Methods for studying metabolism in Drosophila. Methods 68:105–115. doi.org/10.1016/j.ymeth.2014.02.034

Torossian, C. 1968. Recherches Sur La Biologie et l’éthologie De Dolichoderus Quadripunctatus (L) (Hym. Form. Dolichoderidæ). Insectes Sociaux. 15(4):375–88

Volny, V. P., and D. M. Gordon. 2002. Genetic basis for queen-worker dimorphism in a social insect. Proc. Natl. Acad. Sci. USA. 99:6108–6111

Volny, V. P., M. J. Greene, D. M. Gordon, V. P. Volny, M. J. Greene, and D. M. Gordon. 2006. Brood Production and Lineage Discrimination in the Red Harvester Ant (Pogonomyrmex barbatus). Ecology. 87:2194–2200.

Voss, S. H., J. F. McDonald, and C. H. Keith. 1988. Production and abortive development of fire ant trophic eggs. Advances in myrmecology. Pp. 517–534. ed. J.C. Trager. New York : E. J. Brill.

Wardlaw, J. C., and G. W. Elmes. 1995. Trophic eggs laid by fertile Myrmica queens (Hymenoptera: Formicidae). Insectes Soc. 42:303–308. doi.org/10.1007/BF0124042

Wardlaw, J. C., and G. W. Elmes. 1998. Variability in oviposition by workers of six species of Myrmica (hymenoptera, formicidae). Insectes Soc. 45:369–384. doi.org/10.1007/s000400050096

Wei, H., He, X.J., Liao, C.H., Wu, X.B., Jiang, W.J., Zhang, B., Zhou, L.B., Zhang, L.Z., Barron, A.B., Zeng, Z.J. 2019. A Maternal Effect on Queen Production in Honeybees. Current Biology. 29(13):2208-2213.e3. doi.org/10.1016/j.cub.2019.05.059.

Wenseleers, T and F.L.W. Ratnieks. 2006 Comparative analysis supports relatedness theory for worker policing in eusocial Hymenoptera. Am Nat. 168 :E163–79. doi: 10.1086/508619

Wheeler, W. M. 1910. Ants: Their Structure, Development, and Behavior. The Columbia University Press, New York. doi.org/10.1155/1922/80849

Wilson, E. O. 1971. The insect societies. Harvard University Press, Cambridge, MA.

Wilson, E. O. 1976. A social ethogram of the neotropical arboreal ant Zacryptocerus varians (Fr. Smith). Anim. Behav. 24:354–363. doi.org/10.1016/S0003-3472(76)80043-7

Yamada, A., F. Ito, R. Hashim, and K. Eguchi. 2018. Queen polymorphism in Acanthomyrmex careoscrobis Moffett, 1986 in Peninsular Malaysia (Hymenoptera: Formicidae: Myrmicinae), with descriptions of hitherto unknown female castes and males. Asian Myrmecology. 10(e010009):1–19. doi.org/10.20362/am.010009

Yamauchi, K., T. Furukawa, K. Kinomura, H. Takamine, and K. Tsuji. 1991. Secondary Polygyny by Inbred Wingless Sexuals in the Dolichoderine Ant Technomyrmex albipes. Behav. Ecol. Sociobiol. 29:313–319.

Zhu, K., M. Liu, Z. Fu, Z. Zhou, Y. Kong, H. Liang, Z. Lin, J. Luo, H. Zheng, P. Wan, J. Zhang, K. Zen, J. Chen, F. Hu, C. Y. Zhang, J. Ren, and X. Chen. 2017. Plant microRNAs in larval food regulate honeybee caste development. PLoS Genet. 13:1–23. doi.org/10.1371/journal.pgen.1006946

