## Supplementary Tables 1 and 2 for "Trophic eggs affect caste determination in the ant *Pogonomyrmex rugosus*"

E. Genzoni, T. Schwander and L. Keller

### Supplementary material

**Supplementary Table 1.** List of ant species for which previous studies described the presence of trophic eggs (T-eggs). Information is given on whether they are laid by the queen or the workers, or whether no trophic eggs were detected. na indicates that no information is available. [1] Possibly laid by queens at the founding stage, [2] only virgin queens, [F&E] laid at founding and established stages, [F] laid at founding stage, [E] laid at established stage, [F?] or [E?] unknown at founding or established stage, respectively, [U] unknown if laid at founding and/or established stages.

| Subfamily | Species | T-eggs laid by queen | T-eggs laid by worker | References |
|---|---|---|---|---|
| Amblyoponinae | *Amblyopone silvestrii* | No | No | (Masuko 2003) |
| | *Amblyopone sp. (Reclinata group)* | na | Yes | (Ito 1991, 1993) |
| | *Prionopelta kraepelini* | na | Yes | (Masuko 2003) |
| Dolichoderinae | *Dolichoderus quadripunctatus* | na | Yes | (Torossian 1968; Fletcher and Ross 1985) |
| | *Linepithema humile* | na | Yes | (Bartels 1988) |
| | *Technomyrmex albipes* | Yes [F?&E] | Yes | (Yamauchi *et al.* 1991) |
| Ectatomminae | *Ectatomma tuberculatum* | na | Yes | (Hora *et al.* 2007; Azevedo *et al.* 2011) |
| | *Gnamptogenys menadensis* | No | Yes | (Gobin *et al.* 1998, 1999) |
| | *Gnamptogenys costata* | na | Yes | (Gobin *et al.* 1998) |
| | *Gnamptogenys dammemani* | na | Yes | (Gobin *et al.* 1998) |
| | *Gnamptogenys moelleri* | na | Yes | (Gobin *et al.* 1998) |
| Formicinae | *Anoplolepis gracilipes* | na | Yes | (Lee *et al.* 2017) |
| | *Cataglyphis floricola* | na | Yes | (Amor *et al.* 2017) |
| | *Cataglyphis tartessica* | na | Yes | (Amor *et al.* 2017) |
| | *Formica pergandei* | na | Yes | (Hung 1973) |
| | *Lasius niger* | Possible[1] | Yes | (Baroni Urbani 1991; Khila and Abouheif 2008) |
| | *Oecophylla longinoda* | na | Yes | (Hölldobler and Wilson 1983; Fletcher and Ross 1985) |
| | *Plagiolepis pygmaea* | no | Yes | (Passera 1978, 1980; Fletcher and Ross 1985) |
| Myrmeciinae | *Myrmecia forceps* | Yes [U] | Yes | (Freeland 1958) |
| | *Myrmecia gulosa* | na | Yes | (Freeland 1958; Dietemann et al. 2002) |
| Myrmicinae | *Pheidole pallidula* | Yes [2] | No | (Passera 1978; Lorber and Passera 1981; Bartels 1988) |
| | *Solenopsis invicta* | Yes [F&E] | No | (Fletcher and Ross 1985; Voss et al. 1988; Cassill 2002) |
| | *Pogonomyrmex badius* | na | na | (Freeland 1958; Smith et al. 2007) |
| | *Pogonomyrmex barbatus* | Yes [F&E?] | na | (Volny et al. 2006; Smith et al. 2007) |
| | *Pogonomyrmex J lineages* | Yes [U] | na | (Helms Cahan *et al.* 2011) |
| | *Acromyrmex sp.* | No | Yes | (Dijkstra et al. 2005) |
| | *Aphaenogaster (=Novomessor) cockerelli* | na | Yes | (Hölldobler and Carlin 1989; Smith et al. 2008) |
| | *Aphaenogaster rudis* | na | Yes | (Khila and Abouheif 2008, 2010) |
| | *Aphaenogaster subterranea* | na | Yes | (Passera 1978) |

| | | | | |
|---|---|---|---|---|
| | *Atta laevigata* | Yes [F&E?] | Yes | (Passera 1978) |
| | *Leptothorax acervorum* | Unlikely | Yes | (Ito 2005) |
| | *Messor capitatus* | Unlikely | Yes | (Passera 1978; Baroni Urbani 1991) |
| | *Messor semirufus* | Unlikely | Yes | (Baroni Urbani 1991) |
| | *Myrmica americana* | na | Yes | (Khila and Abouheif 2008) |
| | *Temnothorax recedens (Nyl.)* | No | Yes | (Dejean and Passera 1974) |
| | *Zacryptocerus varians (fr. smith)* | na | Yes | (Wilson 1976) |
| | *Acanthomyrmex careoscrobis Moffett* | Yes, by ergatoid queen | Yes | (Yamada et al. 2018) |
| | *Acanthomyrmex ferox* | Yes [F&E] | Yes | (Gobin and Ito 2000) |
| | *Atta sp.* | na | Yes | (Dijkstra et al. 2005) |
| | *Atta sexdens* | Yes [F] | Yes | (Della Lucia *et al.* 1990; Augustin *et al.* 2011) |
| | *Crematogaster smithi Creighton* | Yes [F&E] | Yes | (Heinze et al. 1995, 1999) |
| | *Myrmica rubra* | Yes [F&E] | Yes | (Brian and Rigby 1978; Smeeton 1981; Wardlaw and Elmes 1995, 1998) |
| | *Myrmica ruginodis* | Yes [F&E] | Yes | (Wardlaw and Elmes 1995, 1998) |
| | *Myrmica schencki* | Yes [F&E] | Yes | (Wardlaw and Elmes 1995, 1998) |
| | *Myrmica sulcinodis* | Yes [F&E] | Yes | (Wardlaw and Elmes 1995, 1998) |
| Nothomyrmeciinae | *Nothomyrmecia macrops* | na | Yes | (Taylor 1978) |
| Ponerinae | *Hypoponera eduardi* | na | No | (Choe 1988) |
| | *Odontomachus haematodes* | na | No | (Colombel 1972) |
| | *Rhytidoponera purpurea* | na | na | (Haskins and Whelden 1965) |
| | *Pachycondyla apicalis* | No | Yes | (Dietemann and Peeters 2000) |
| | *Neoponera villosa* | na | Yes | (CamargoMathias and Caetano 1995) |
| | *Pachycondyla krugeri* | na | Yes | (Gobin *et al.* 1998) |
| | *Paraponera clavata* | Unlikely | Yes | (Peeters 2017) |
| | *Diacamma rugosum* | No queen | na | (Wheeler and Chapman 1922) |

**Supplementary Table 2**. Number of worker and queen pupae that developed in recipient colonies without trophic eggs (W) or with three trophic eggs (3T). Empty cells mean that no pupae developed in this recipient colony.

| Sample size | 50 | | 61 | | 66 | | 69 | | 75 | | 81 | | 117 | | 120 | | 125 | | 130 | | 150 | | 173 | |
|---|---|---|---|---|---|---|---|---|---|---|---|---|---|---|---|---|---|---|---|---|---|---|---|---|
| | W | 3T | W | 3T | W | 3T | W | 3T | W | 3T | W | 3T | W | 3T | W | 3T | W | 3T | W | 3T | W | 3T | W | 3T |
| **worker** | | 5 | | 12 | 1 | 2 | 1 | | | 4 | | 1 | | 1 | 1 | 2 | | 1 | 1 | | 1 | 2 | | |
| **gyne** | 2 | 3 | 3 | | 1 | | 1 | 1 | 2 | 1 | 1 | | | | 1 | | | | | | 2 | 1 | 2 | |

| Sample size | 193 | | 215 | | 235 | | 286 | | 303 | | 316 | | 338 | | 12B | | 59A | | 99A | | Total | |
|---|---|---|---|---|---|---|---|---|---|---|---|---|---|---|---|---|---|---|---|---|---|---|
| | W | 3T | W | 3T | W | 3T | W | 3T | W | 3T | W | 3T | W | 3T | W | 3T | W | 3T | W | 3T | W | 3T |
| **worker** | | | | | | 1 | | 4 | 1 | | | 1 | | 2 | | 5 | | 3 | | | 6 | 46 |
| **gyne** | 3 | 3 | | 2 | | | 2 | | | 1 | 2 | 1 | 1 | 1 | 4 | | 2 | 1 | | 2 | 29 | 17 |
